# Reprogramming the cells secretory machinery: A cystic fibrosis rescue

**DOI:** 10.1101/2023.12.09.570886

**Authors:** Won Jin Cho, Bhanu P. Jena

## Abstract

Cystic fibrosis (CF) is a genetic disorder resulting in the secretion of highly viscous mucus in the airways, leading to lung infection, respiratory failure, morbidity and mortality. CF is attributed to mutation in the CF Transmembrane Conductance Regulator (CFTR) gene that codes for a chloride transporting channel at the cell plasma membrane. More than 2,000 mutations in the CF gene have been identified; however, the ΔF508 CFTR is the most common, accounting for approximately 70% of all CFTR mutations. Our earlier studies demonstrate the CFTR protein to be among the nearly 30 proteins constituting the porosome secretory machinery at the plasma membrane in human airway epithelial mucous-secreting cells. Additionally, our recent studies show that stimulated human airway epithelial cells pre-exposed to CFTR inhibitors result in loss of mucus secretion, suggesting the involvement of CFTR in porosome-mediated mucus secretion. To further test the hypothesis that CFTR is involved in porosome-mediated mucus secretion in the human airways, and to develop a therapeutic approach to overcome this defect in ΔF508 CFTR human bronchial epithelial cells, the current study was undertaken. Mass spectrometry and Western Blot analysis of porosomes isolated from WT-CFTR Human Bronchial Epithelial (HBE) Cells and ΔF508-CFTR CF HBE cells, demonstrate a varying loss or gain of several porosome proteins, including a decrease in the t-SNARE protein SNAP-23 and undetectable levels of the Ras GTPase activating like-protein IQGAP1 in the ΔF508-CFTR CF cells. This suggested that mutation in porosome-associated CFTR protein additionally affects other proteins within the porosome secretory machinery, negatively impacting mucus secretion. Hence, to ameliorate defects in mucus secretion in CF, the reconstitution of functional porosomes obtained from WT-CFTR HBE cells into the plasma membrane of ΔF508-CFTR mutant cells was performed. Results from the study demonstrate that porosome reconstitution rescues mucus secretion approximately four-fold more effectively than the currently available CF drugs Tezacaftor and Ivacaftor.

## 1. INTRODUCTION

### 1.1 Lung Mucus in Cystic Fibrosis

Human airway epithelium is coated with a thin film of mucus, composed primarily of mucin MUC5AC and MUC5B [1]. Mucus is continually moved via ciliary action and cleared, assisting both in lubrication and keeping the airway moist, clean and free from infection. Mucus hydration has been recognized as a major issue in cystic fibrosis (CF), resulting in increased mucin viscosity and therefore the inability of the cilia to propel mucin, leading to its stagnation and infection. Studies report that in CF patients, there is loss of MUC5AC and MUC5B mucin in sputum [2]. Sputum from patients with CF, show a 70% decrease in MUC5B and a 93% decrease in MUC5AC in their sputum compared to subjects without any lung disease (*P* < 0.005). Our recent studies in human airway epithelial cell line Calu-3, also demonstrate loss of both chloride and mucus secretion [3] following exposure to the thiazolidinone CFTR inhibitor 172 and the hydrazide CFTR inhibitor GlyH101. Furthermore, since proteome of the porosome secretory machinery in Calu-3 cells is composed of the CFTR protein [4], suggests CFTR involvement in porosome-mediated mucus secretion in the human airway epithelia.

### 1.2 Porosome-Mediated Secretion

In mammalian cells, secretion [5,6] universally involves the docking, swelling and transient fusion of secretory vesicles at cup-shaped lipoprotein structures called porosomes [4–34] at the plasma membrane. Porosome-mediated secretion, enables the regulated and measured release of intra-vesicular products from cells. Secretory vesicles undergo swelling involving the regulated rapid transport of ions and water via aquaporin water channels [35–41], during SNARE-mediated fusion of secretory vesicle at the porosome [42–50]. Secretory vesicle membrane fuse at the base of the porosome complex, to expel intra-vesicular contents from the cell. Vesicle swelling result in hydration and build-up of intra-vesicular pressure, required for content expulsion during cell secretion. Greater the vesicular pressure, more the contents that are released. This precisely regulated mechanism of cell secretion, helps retains the integrity of the secretory vesicle membrane and the cell plasma membrane [51–53]. The cup-shaped porosome complex range in size from 15 nm in neurons and astrocytes, to 100-180 nm in endocrine and exocrine cells. In Calu-3 cells, porosomes measure approximately 100 nm, similar to insulin-secreting beta cells of the endocrine pancreas.

### 1.3 Functional Reconstitution of the Porosome Complex in Lipid Membranes

In the past two decades, to test whether the immuno-isolated porosome preparations are intact and functional, purified porosomes obtained from various cell types have been functionally reconstituted into artificial lipid membranes and live cells. Porosomes have been isolated from acinar cells of the exocrine pancreas [15], from insulin-secreting beta cells of the endocrine pancreas [28] and from brain neurons [11], their chemistry determined using mass spectrometry and Western blot analysis [7,11,12,14–18,26,27], and they have been functionally reconstituted [28]. Functional reconstitution of the porosome secretory machinery into artificial lipid membranes, has been accomplished using established electrophysiological bilayer setups (EPC9) having a 200 µm diameter lipid bilayer membrane separating a *cis* and a *trans* compartment. Lipid bilayers were prepared using phosphatidylethanolamine (PE) and phosphatidylcholine (PC) obtained from Avanti Lipids (Alabaster, AL). A suspension of PE:PC in 7:3 ratio was used to establish a stable bilayer in a 200 µm in diameter hole, by brushing the lipid mixture. The bilayer membranes were established while holding the membrane potential at 0 mV, and once a bilayer was established and demonstrated to be in the capacitance limits for a stable bilayer membrane, the voltage was switched to −60 mV. An established stable bilayer exhibited a capacitance typically between 100 and 250 pF. Thereafter, once a baseline current was established, isolated porosomes were reconstituted into the bilayer membrane by brushing in isolated porosome preparations from a specific cell type. Isolated secretory vesicles depending on the cell type, such as synaptic vesicles from neurons or zymogen granules from exocrine pancreas, were then added to the *cis* compartment of the bilayer chamber, and the capacitance and conductance at the bilayer is measured that provides vesicle docking and fusion events. To biochemically monitor the release of intra-vesicular contents into the *trans* compartment, aliquots from the *cis* and *trans* chamber were collected at different time intervals, followed by Western Blot analysis. Data obtained from the electrical activity of the porosome-reconstituted membrane and the immunoblot analysis of the transport of vesicular contents from the *cis* to the *trans* compartments of the bilayer chambers, demonstrated whether the membrane-reconstituted porosomes were indeed functional.

This *in vitro* porosome-mediated setup, now provides a tool to understand the role of various molecules that could modulate secretory vesicle docking and fusion at the porosome complex. For example, it has been demonstrated that calcium is required for fusion of opposing bilayers. Similarly, GTP is required for secretory vesicle swelling and content release during cell secretion. In reconstituted porosome complex obtained from exocrine pancreas, chloride channel activity is present and is found to be critical to its activity, since the chloride channel blocker DIDS inhibits reconstituted porosome-mediated intra-vesicular content release.

### 1.4 Functional Reconstitution of the Porosome Complex in Live Cells

Functional porosome reconstitution in live cells have also been well established. For example, in Min6 mouse insulinoma cells, insulin-secreting porosome reconstitution exhibit glucose-stimulated insulin secretion. Isolated Min6 porosomes reconstituted into live Min6 cells exhibit an increase in potency and efficacy of glucose-stimulated insulin release within 1 h following reconstitution [28]. This glucose-stimulated insulin release is sustained in these reconstituted cells beyond a 48h period, demonstrating functional reconstitution of porosomes into live cells [28].

### 1.5 Porosome-Mediated Mucus Secretion and CF Therapy

Given the nearly 30-years of progress made in our understanding of the secretory process in cells following the porosome discovery, the current study was undertaken to determine **a)** the impact of ΔF508 CFTR mutation on the proteome of the mucin-secreting porosome complex in HBE cells, and **b)** the effect of reconstitution of normal functional porosomes isolated from WT-CFTR HBE cells into ΔF508 CFTR HBE CF cells. The proteome of the porosome in WT-CFTR and ΔF508-CFTR HBE CF cells was first studied to determine any change in the porosome secretory complex as a result of the ΔF508 mutation in the CFTR. Mass spectrometry and Western Blot analysis of porosomes isolated from WT-CFTR HBE cells and ΔF508-CFTR HBE CF cells, demonstrate a decrease in the t-SNARE protein SNAP-23 and undetectable levels of the ras GTPase activating like-protein IQGAP1 in ΔF508 cells. This suggested that mutation in porosome-associated CFTR protein additionally affects other proteins within the porosome secretory machinery, that would negatively impacting mucus secretion. These results further imply that just correcting or modulating the function of the mutated CFTR protein in CF would be inadequate in overcoming the disease. Rather, all proteins within the porosome complex that are impacted by different CFTR mutations need to be addressed. This would be a near impossible task, since the variability in changes to other proteins within the porosome machinery as a result of the nearly 2,000 or so CFTR mutations would be complex. To overcome this issue and to be able to rescue altered mucus secretion in all CF mutations, the reconstitution of functional porosomes obtained from WT-CFTR HBE cells into the plasma membrane of ΔF508-CFTR mutant cells, were performed. Results from the study demonstrate

that porosome reconstitution rescues mucus secretion four-fold more effectively than two of the currently available CF drugs namely Tezacaftor and Ivacaftor.

## 2. MATERIALS AND METHODS

### 2.1 Human Bronchial Epithelial Cell Line WT-CFTR and ΔF508-CFTR Cultures

WT-CFTR Human Bronchial Epithelial (HBE) cell line (CFBE41o-6.2) and experimental ΔF508-CFTR HBE CF cell line (CFBE41o) a ΔF508del (-/-) homozygous, was obtained from Sigma (Temecula, CA 92590, USA). Cells were stored, handled and cultured using established published procedures and protocols outlined in the data sheet. Cells were cultured using Fibronectin/Collagen/BSA ECM mixture-coated petri dishes (10µg/mL Human Fibronectin, Sigma Cat. No. F2006; 100µg/mL BSA, Sigma Cat.No.126575; 30µg/mL PureCol, Sigma Cat No. 5006; α-MEM Medium, Sigma Cat. No. M2279) with α-MEM Medium supplemented with 2mM L-Glutamine (Gibco Cat. No. 25030-081, Grand Island, NY 14072, USA); 10% fetal bovine serum (Sigma Cat. No. ES-009-B); and 100 U/ml Penicillin and 100 µg/ml Streptomycin (Gibco Cat. No. 15140-122). Cultures were incubated at 37°C in 95% air/5% CO2 atmosphere.

### 2.2 Differentiated Human Bronchial Epithelial 3D Cell Cultures

Air liquid interface (ALI) differentiated WT-CFTR and ΔF508-CFTR CF HBE cell cultures mimicking normal lung physiology, were established using a minor modification of our published procedure [3,4]. These ALI 3D cultures respond to CFTR inhibitors [3] and the currently available CF corrector/modulator drugs Tezacaftor and Ivacaftor, as presented in the present study. To establish ALI 3D cultures, cells were seeded in sterile ECM mixture-coated transwell inserts of 12 mm diameter with 0.4 µm pore size (Corning Cat. No. Costar 3460, Kennebunk. ME 04043 USA) with a seeding density of 2 x 10^5^ cells/insert. Media of apical and basolateral region of transwell were changed every other day, 500µL on the apical side and 1mL on the basolateral side. At confluency on day 7, cells were raised to ALI condition without apical medium. On day 21 after switching to ALI, cultures were treated with either Tezacaftor (5µM, Selleck Chemicals Cat. No. S7059, Houston, TX 77014 USA), Ivacaftor (5µM, Selleck Chemicals Cat. No. S1144) or immuno-isolated porosomes (1µg/mL).

### 2.3 Porosome Isolation and Reconstitution

WT-CFTR HBE cell line (CFBE41o-6.2) and experimental ΔF508-CFTR HBE CF cell line (CFBE41o) a ΔF508del (-/-) homozygous were used to isolate porosomes for proteome analysis and reconstitution. SNAP-23 specific antibody (Abcam Cat. No. AB3340, Cambridge, UK) was used to immunoisolate porosomes from solubilized cells. Protein in all fractions was estimated using BCA Protein assay Kit (ThermoFisher Cat. No. 23227, Rockford, IL 61101, USA). SNAP-23 specific antibody-crosslinked to protein A/G Magnetic-agarose (ThermoFisher Cat. No. 78609) was used. To reduce the antibody contamination in eluted protein solution, the antibody was chemically crosslinked to the agarose-magnetic beads. Briefly, the beads were resuspended in dilution buffer (1:1 ratio, 1mg/mL BSA in PBS) for 10 min at 4°C, centrifuged for 1 min at 14,000 rpm in a bench top centrifuge, and the supernatant aspirated. SNAP-23 antibody (1µg/mL) in dilution buffer was added to the beads at 1:1 ratio and mixed gently for 1hr at 4°C. The beads were then washed twice with 10 volumes of dilution buffer. Dimethyl pimelimidate solution (DMP, 13mg/ml, Sigma Cat. No. D8388) in wash buffer (0.2 M triethanolamine in PBS, Sigma Cat. No. 90279) was added to the SNAP23 antibody conjugated beads (1:1 ratio) and resuspended for 30 min at room temperature (RT). The beads were then washed with wash buffer three time. (30 min/wash at RT), and resuspended in quenching buffer (50mM ethanolamine in PBS, Sigma Cat. No. E0135) for 5 min at RT and washed with PBS twice. To remove excess unlinked antibody, the beads were washed with 1 M glycine pH 3, twice (10min/wash at RT). Prior to use for immunoprecipitation, the beads were washed in PBS-TWEEN buffer three times. Cells were solubilized in Triton/Lubrol solubilization buffer (0.5% Lubrol; 1 mM benzamidine; 5 mM ATP; 5 mM EDTA; 0.5%Triton X-100, in PBS), supplemented with protease inhibitor mix (Sigma, St. Louis, MO). SNAP-23 antibody-crosslinked to the protein A/G Magnetic-agarose was incubated with the solubilized cell lysates for 16h at 4°C followed by washing with wash buffer (500 mM NaCl, 10 mM TRIS, 2 mM EDTA, pH 7.5).

The immune-isolated porosomes associated with the immuno-agarose beads were dissociated and eluted using pH 3.0 PBS solution, and the eluted sample was immediately returned to neutral pH prior to use in mass spectrometry, Western Blot analysis and reconstitution. For porosome reconstitution, ΔF508-CFTR CF HBE cells were exposed to 1µg/ml porosomes isolated from WT-CFTR HBE cells.

### 2.4 Mass Spectrometry on Isolated Porosomes

Porosomes isolated from expanded WT-CFTR HBE cells (CFBE41o-6.2) and experimental ΔF508-CFTR CF HBE cells (CFBE41o) were subjected to mass spectrometry. Samples were digested with trypsin and analyzed on an Orbitrap Eclipse MS system. Data were analyzed in Proteome Discoverer 2.4 using Sequest and Percolator algorithms. Values obtained represent multiple consensus-based quantitation.

### 2.5 Western Blot Analysis on Isolated Porosomes

Total cell homogenates (TH) and porosomes immuno-isolated (IP) from expanded WT-CFTR HBE cell culture (CFBE41o-6.2) and experimental ΔF508-CFTR CF HBE cell (CFBE41o) cultures were subjected to SDS-PAGE and Western Blot analysis. 20µg of proteins (TH) and 10µL of isolated porosomes in Laemmli buffer were resolved in a 12.5% SDS-PAGE, followed by electrotransfer to 0.2-mm nitrocellulose (NC) membrane. The NC membrane was incubated for 1 hour at RT in blocking buffer (5% nonfat milk in PBS [pH 7.4] containing 0.1% Triton X-100 and 0.02% NaN3) and immunoblotted for 2 hours at RT with antibodies to CFTR (1:1000, Cell signaling Technology Cat. No. 78335,), SNAP-23 (1:1000, Abcam Cat. No. AB3340) and GAPDH (1:3,000, Santa Cruz Cat. No. sc-25778, Dallas, TX 75200, USA). Anti-rabbit horseradish peroxidase secondary antibody conjugates were used (1:5,000, Cell Signaling Technology Cat. No. 7074), followed by the NC membrane developed using Western Lightning Plus-ECL (PerkinElmer, Waltham, MA 02451. USA) and imaged using ChemiDoc XRT+ image system (Bio-Rad, Richmond, CA94806, USA).

### 2.6 ELISA

At different time point (1day, 2day) following treatment with Tezacaftor, Ivacaftor or isolated porosomes, the apical surface, basal surface, or both, of each ALI culture, was gently washed and aspirated using 500uL fresh PBS. These washes were stored at 4°C for ELISA assays for MUC5AC and MUC5B. To quantify the mucin secretion, 100uL of each wash was added to the ELISA plate. MUC5AC ELISA Kit (MyBioSource Cat. No. MBS701926, San Diego, CA 92195, USA) and MUC5B ELISA Kit (MyBioSource Cat. No. MBS2024599) were used according to the manufacturer’s instructions. Optical density was determined at 450nm using a BioTek Synergy HT microplate reader (BioTek, Winooski, VT 05404, USA), and the mucins quantified.

### 2.7 Light Microscopy

WT-CFTR HBE cells (CFBE41o-6.2) and experimental ΔF508-CFTR CF HBE cells (CFBE41o) in culture, were imaged at different intervals to determine cell health and viability using a Leica DM IL microscope (Leica Microsystems, Switzerland).

## 3. RESULTS AND DISCUSSION

ALI differentiated WT-CFTR and ΔF508-CFTR CF HBE 3D cultures []3,4] respond to CFTR inhibitors and CF corrector/modulator drugs Tezacaftor and Ivacaftor, as presented in the study. Both WT-CFTR HBE cell cultures and the ΔF508-CFTR CF HBE cell cultures were regularly imaged at different intervals, and determined to be health [Figure 1]. This ALI 3D model closely mimicking the physiological state of human airway epithelium, is of great help in assessing CF treatments and therapy prior to human clinical trials. Western blot analysis of porosomes isolated from WT-CFTR HBE expanded cell cultures (CFBE41o-6.2) and ΔF508-CFTR CF HBE expanded cell cultures (CFBE41o), demonstrated a loss of the t-SNARE protein SNAP-23 in the ΔF508-CFTR cells [Figure 2]. Similarly, mass spectrometry of porosomes isolated from WT-CFTR Human Bronchial Epithelial (HBE) Cells and ΔF508-CFTR CF HBE cells, demonstrate a varying loss or gain of several porosome proteins, including undetectable levels of the Ras GTPase activating like-protein IQGAP1 in the ΔF508-CFTR CF cells [Table I]. These studies demonstrate for the first time that mutation in CFTR impacts other proteins within the porosome secretory machinery besides CFTR. Therefore, expression of just the normal CFTR protein in treating CF would be inadequate in treating the disease. Furthermore, in a multi-protein complex such as the mucus-secreting porosome machinery composed of around 34 proteins, it would be nearly impossible to replace the mutated CFTR protein. It would be even harder to determine and correct changes in other porosome proteins in the defective porosome complex in CF. Due to this issue, it is likely that earlier attempts in using gene therapy, has not been very effective in treating CF. Therefore, current successful CF treatments use modulator products focused on correcting to various degrees (up to 13-14%), the malfunctioning CFTR protein in CF patients. However, none of today’s CF therapies have considered treating the disease as a defect in the mucus secretory machinery of the cell. Although the disease negatively impacts multiple systems in the body, the effects on the respiratory system is the primary contributor to the high morbidity and mortality in CF.

**Table I.**
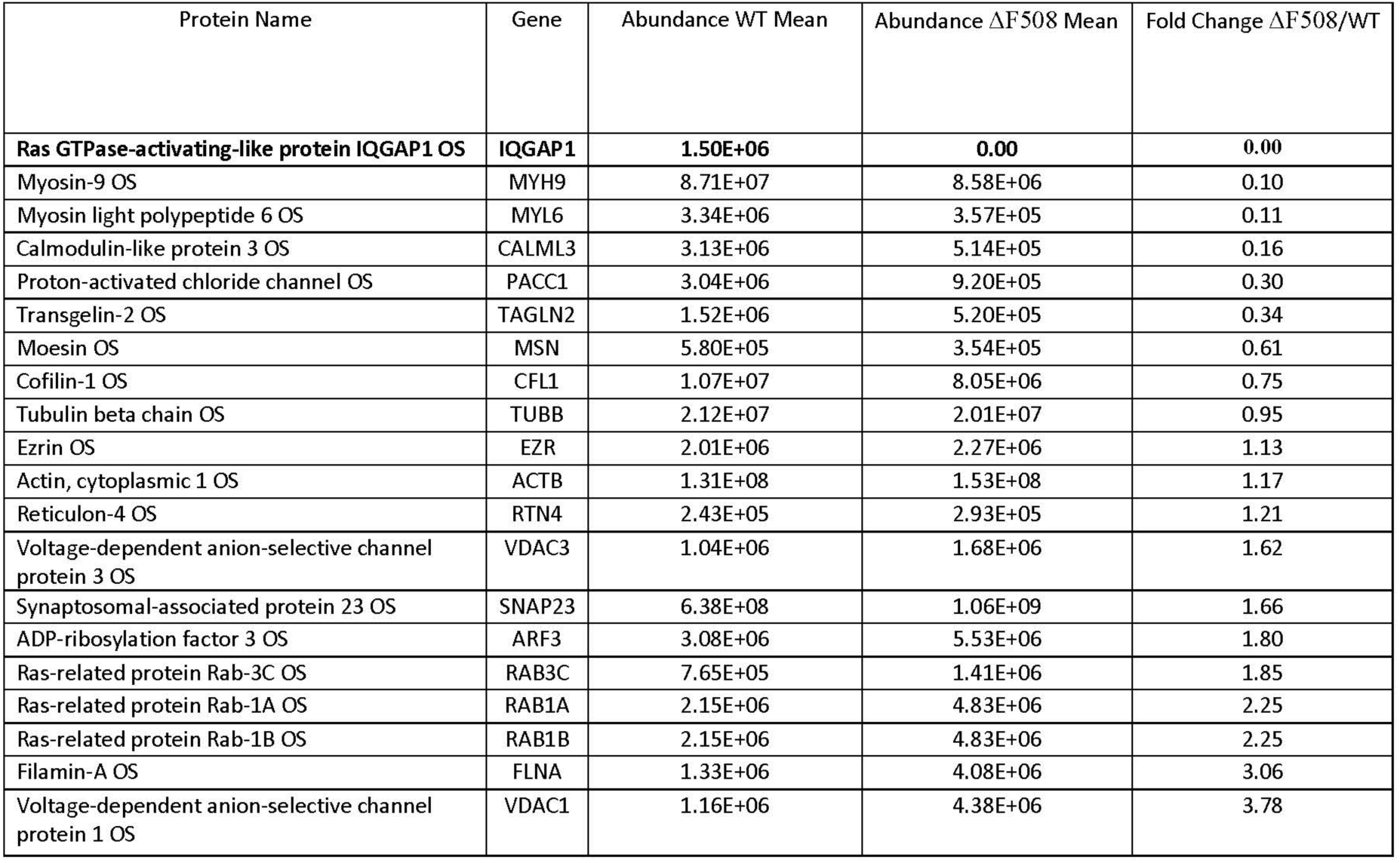
Mass spectrometry on porosome isolated from WT-CFTR HBE cells and ΔF508-CFTR CF cells. Note, that in the porosome complex isolated from the ΔF508-CFTR CF cells, there is a loss or gain of various porosome proteins to varying degrees, including the absence of Ras GTPase-activating-like protein IQGAP1. This is a partial list of the porosome proteins present in HBE cells.

| Protein Name | Gene | Abundance WT Mean | Abundance $\Delta F508$ Mean | Fold Change $\Delta F508$/WT |
| --- | --- | --- | --- | --- |
| **Ras GTPase-activating-like protein IQGAP1 OS** | **IQGAP1** | **1.50E+06** | **0.00** | **0.00** |
| Myosin-9 OS | MYH9 | 8.71E+07 | 8.58E+06 | 0.10 |
| Myosin light polypeptide 6 OS | MYL6 | 3.34E+06 | 3.57E+05 | 0.11 |
| Calmodulin-like protein 3 OS | CALML3 | 3.13E+06 | 5.14E+05 | 0.16 |
| Proton-activated chloride channel OS | PACC1 | 3.04E+06 | 9.20E+05 | 0.30 |
| Transgelin-2 OS | TAGLN2 | 1.52E+06 | 5.20E+05 | 0.34 |
| Moesin OS | MSN | 5.80E+05 | 3.54E+05 | 0.61 |
| Cofilin-1 OS | CFL1 | 1.07E+07 | 8.05E+06 | 0.75 |
| Tubulin beta chain OS | TUBB | 2.12E+07 | 2.01E+07 | 0.95 |
| Ezrin OS | EZR | 2.01E+06 | 2.27E+06 | 1.13 |
| Actin, cytoplasmic 1 OS | ACTB | 1.31E+08 | 1.53E+08 | 1.17 |
| Reticulon-4 OS | RTN4 | 2.43E+05 | 2.93E+05 | 1.21 |
| Voltage-dependent anion-selective channel protein 3 OS | VDAC3 | 1.04E+06 | 1.68E+06 | 1.62 |
| Synaptosomal-associated protein 23 OS | SNAP23 | 6.38E+08 | 1.06E+09 | 1.66 |
| ADP-ribosylation factor 3 OS | ARF3 | 3.08E+06 | 5.53E+06 | 1.80 |
| Ras-related protein Rab-3C OS | RAB3C | 7.65E+05 | 1.41E+06 | 1.85 |
| Ras-related protein Rab-1A OS | RAB1A | 2.15E+06 | 4.83E+06 | 2.25 |
| Ras-related protein Rab-1B OS | RAB1B | 2.15E+06 | 4.83E+06 | 2.25 |
| Filamin-A OS | FLNA | 1.33E+06 | 4.08E+06 | 3.06 |
| Voltage-dependent anion-selective channel protein 1 OS | VDAC1 | 1.16E+06 | 4.38E+06 | 3.78 |

**Figure 1.**
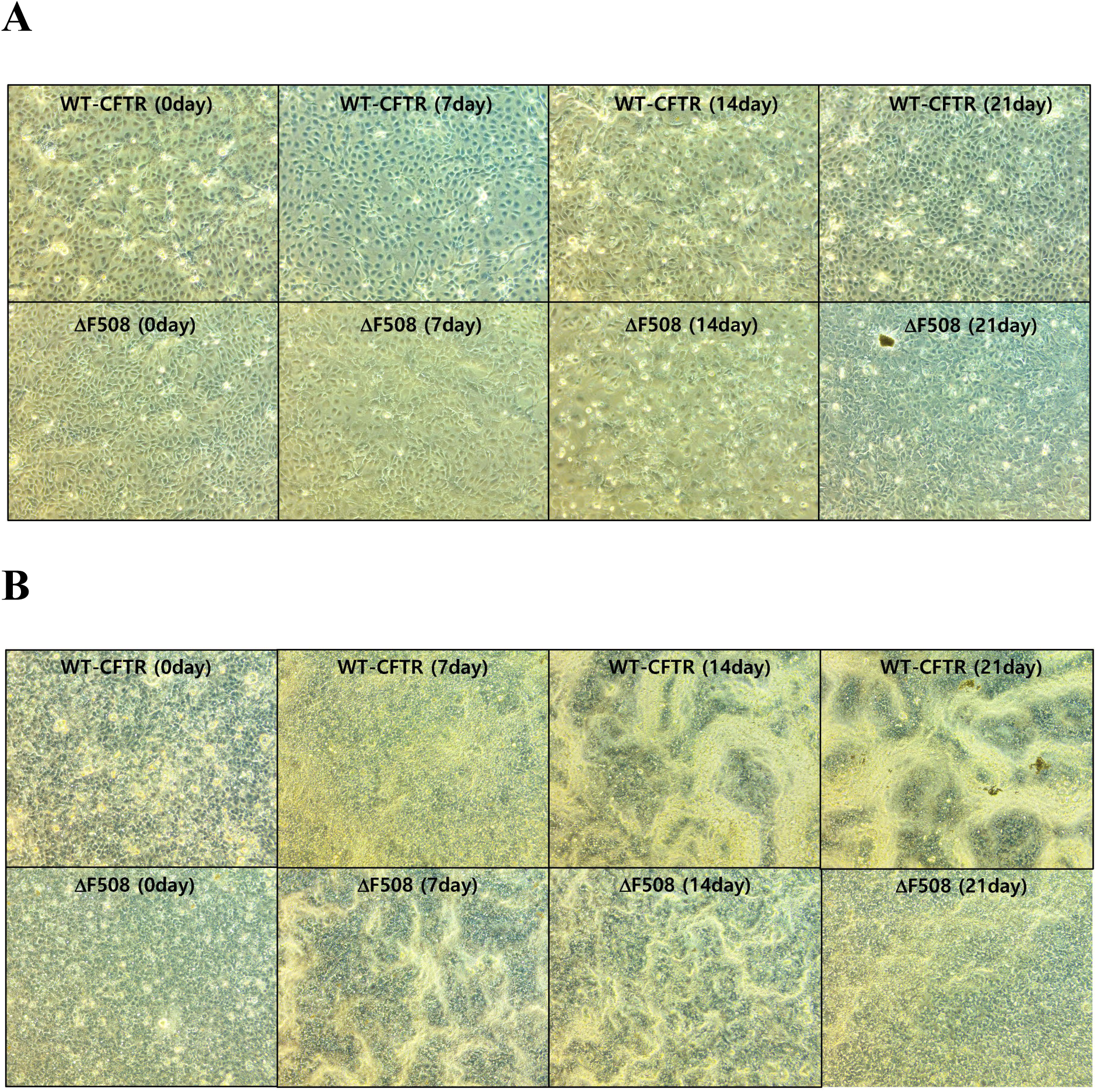
Light microscopy phase images of WT-CFTR HBE cells and ΔF508-CFTR HBE cells at different days following seeding and expansion in petri dishes liquid cultures **(A)** and in air liquid interface (ALI) transwell cultures for differentiation **(B)**. At confluency on day 7, cells were raised to ALI condition without apical medium. Images of ALI cultures **(B)** at day 14 and day 21 are presented. On day 21 after switching to ALI, the ΔF508-CFTR HBE cell cultures were treated with drug or reconstituted using isolated porosome complexes derived from expanded petri dish cultures **(A)**. Note in the ALI cultures in **(B)**, there is greater amounts of mucin in the WT-CFTR cultures compared to the ΔF508-CFTR HBE cells. Throughout the experiment, cultures were devoid of contamination and appear healthy.

**Figure 2.**
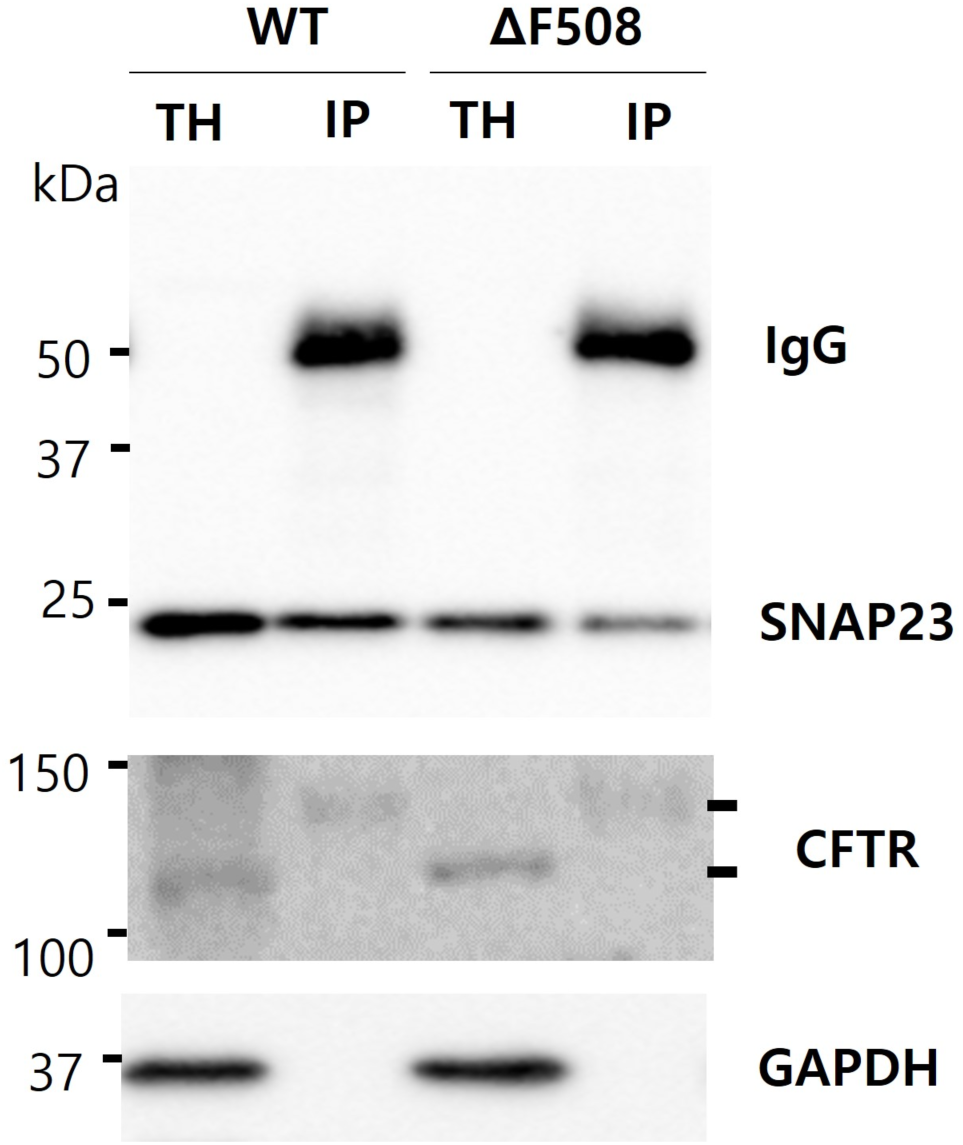
Porosome isolated from WT-CFTR HBE cells and ΔF508-CFTR CF HBE cells using SNAP 23 antibody demonstrate a decrease in the presence of SNAP-23 protein both in total homogenate (TH) and in immune-isolated (IP) porosomes from ΔF508-CFTR CF HBE cells. These results suggest that altered CFTR protein within the porosome complex as a consequence of mutation in the CFTR gene, alters another protein within the porosome complex, altering porosome function. However, the amount of the CFTR protein in the porosome of ΔF508-CFTR CF HBE cells do not appear to change and is present in both the WT and ΔF508 cells in the high molecular glycosylated state. In TH of WT and ΔF508 cells, both the non-glycosylated low molecular weight CFTR and the glycosylated higher molecular weight CFTR forms are present.

To address the issue of multi-protein malfunction in CF in addition to correcting the CFTR protein, and to be able to treat all forms of CFTR mutations, the porosome reconstitution therapy [Figure 3] was performed on ALI differentiated 3D cultures of ΔF508-CFTR HBE CF cells. Porosome reconstitution in ΔF508-CFTR HBE CF cells was able to greatly increase MUC5B and MUC5AC secretions. Two currently available CF drugs, namely Tezacaftor and Lvacaftor, were also used in the study to compare effectiveness in restoring normal MUC5B and MUC5AC secretions. Results from the study [Figure 4, 5] show that on average, porosome reconstitution was able to restore by >40% the secretion of MUC5B in ΔF508-CFTR cells. Similarly, in ΔF508-CFTR CF cells, a 9% and 11% increase in MUC5AC secretion is observed in the presence of Lvacaftor and Tezacaftor respectively, only on day 2 following exposure to the drugs, while porosome-reconstitution in the same period demonstrated a significant 43.8% increase (P<0.05), a near four-fold greater than the two CF drugs. These results show great promise of the porosome-reconstitution therapy in treating CF including all forms of CFTR mutations.

**Figure 3.**
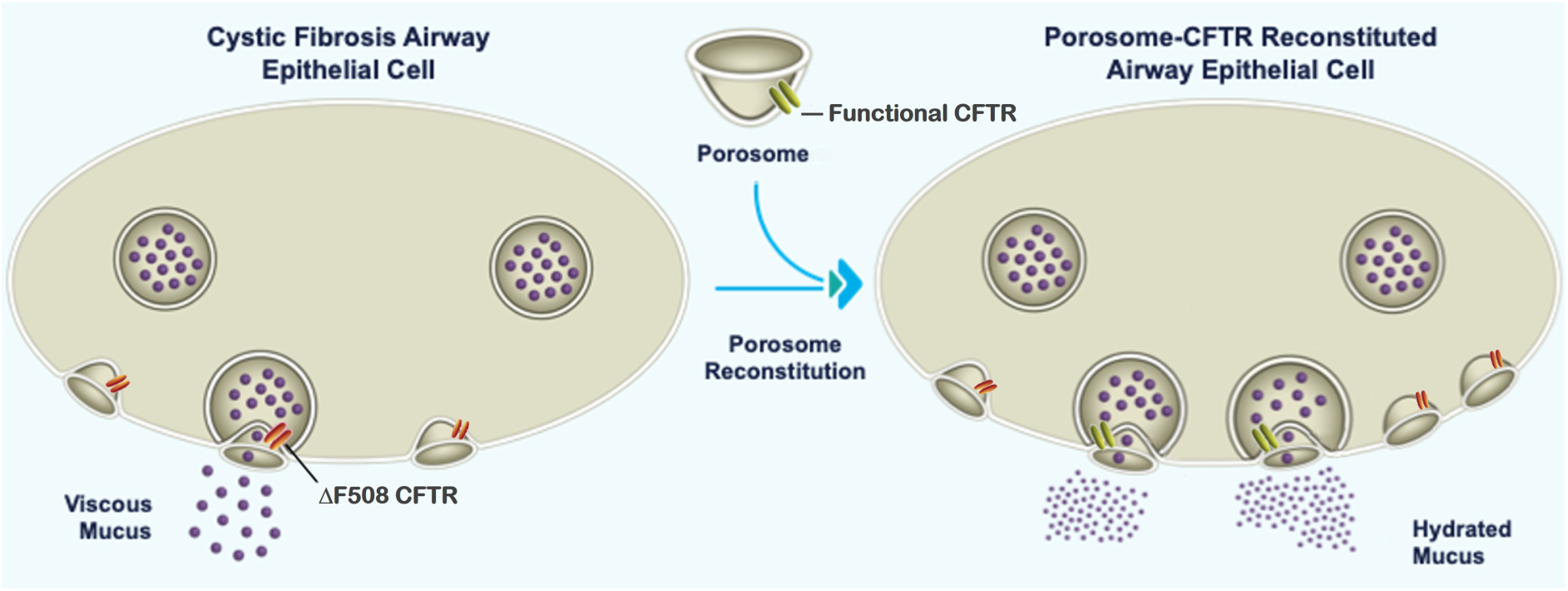
Schematic representation of porosome-reconstitution therapy for cystic fibrosis (CF). Functional normal CFTR porosomes obtained from WT-CFTR human bronchial epithelial cells, will rescues CF by normalizing mucus secretion.

**Figure 4.**
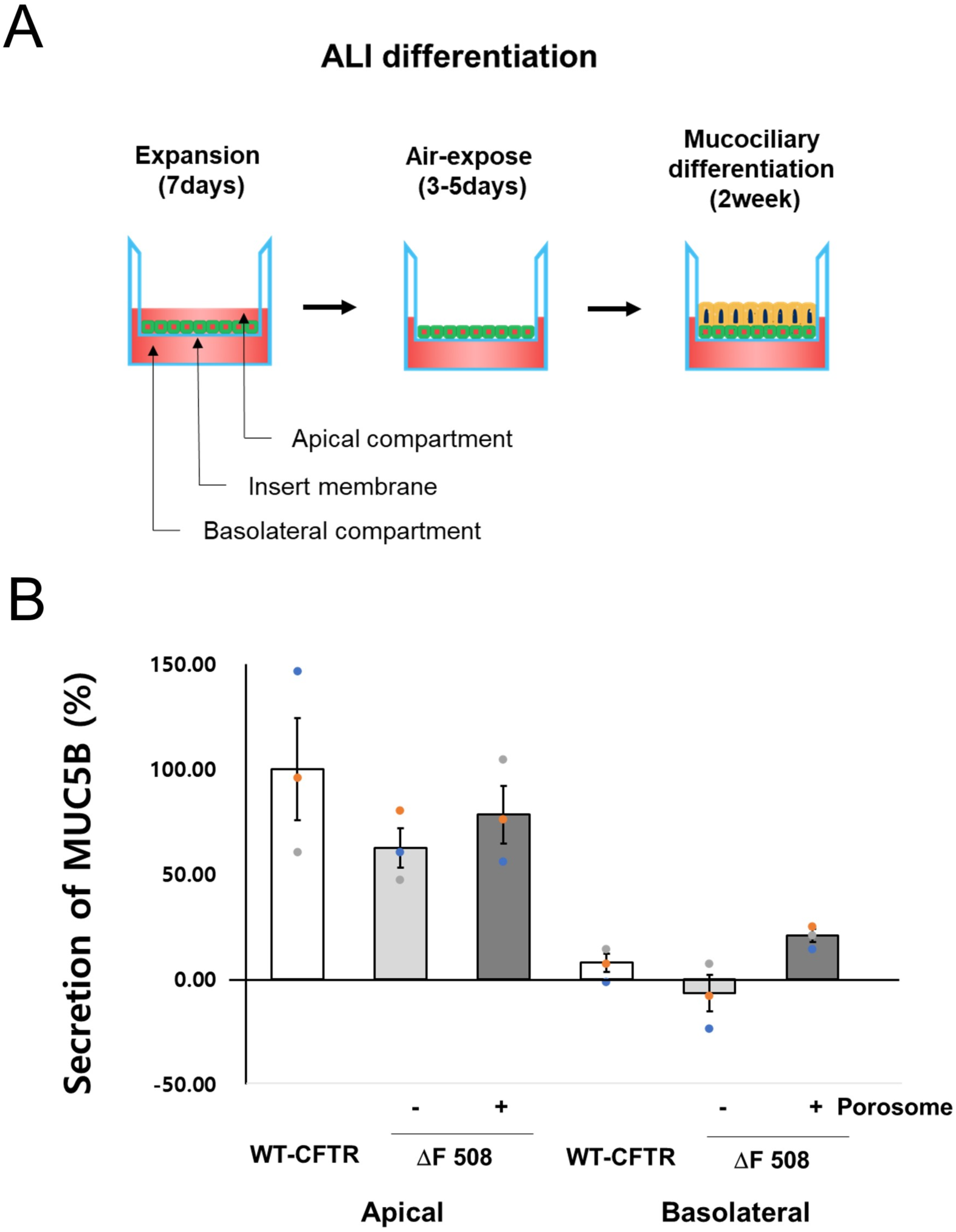
Porosome reconstitution therapy enhances MUC5B secretion in air liquid interface (ALI) differentiated ΔF508-CFTR HBE cell culture. (A) Schematic drawing showing the establishment of AIL cultures. (B) MUC5B secretion is increased in ALI differentiated ΔF508-CFTR HBE cells following porosome reconstitution in day 1. Cells were exposed to 1 µg/ml of isolated porosome complex. Although at the apical end there is clearly an increase in MUC5B secretion in ΔF508-CFTR HBE cells, at the basolateral end, porosome reconstitution demonstrates a greater increase in MUC5B secretion in the in ΔF508-CFTR HBE cells (*P<0.05).

**Figure 5.**
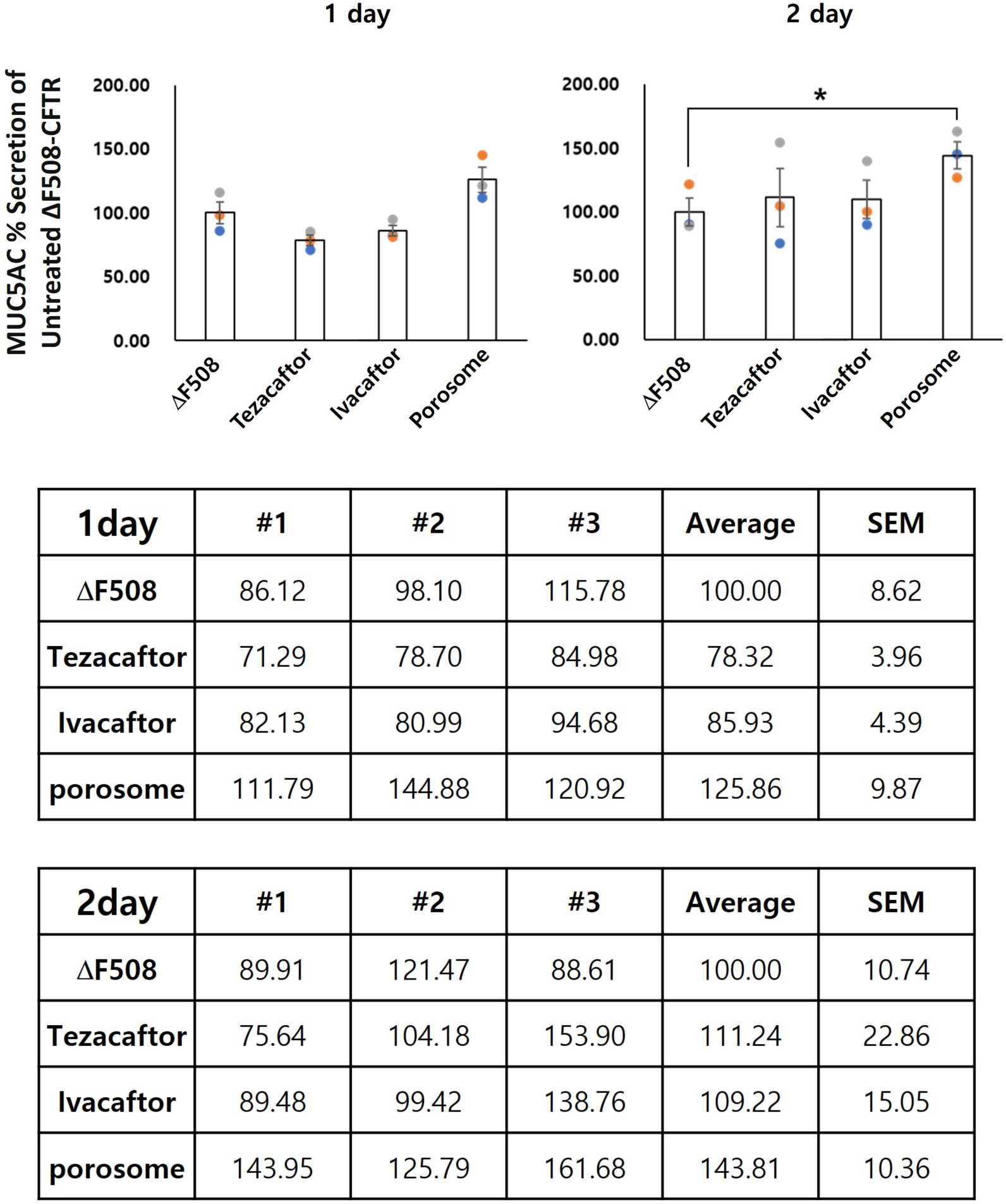
Porosome reconstitution therapy is significantly more efficacious in restoring normal mucin secretion in ΔF508-CFTR human bronchial epithelial cells than Tezacaftor, or Lvacaftor. Data presented as percent of secretion in comparison to untreated ΔF508-CFTR mucociliary cells. MUC5AC secretion from ALI differentiated ΔF508-CFTR human bronchial epithelial cells reconstituted with normal porosomes or exposed to Ivacaftor or Tezacaftor, demonstrates that the porosome reconstitution therapy is significantly more efficacious in restoring MUC5AC secretion in the ΔF508-CFTR CF HBE cells compared to both drugs. Note that while on average, a 9-11% increase in MUC5AC secretion is observed in the presence of Ivacaftor and Tezacaftor respectively on day 2 following exposure of the drugs, only in the porosome-reconstituted cells, a significant Increase in secretion (44%) is observed (*P<0.05). Raw data in percent of MUC5AC secreted in day 1 and 2 following drug exposure or porosome reconstitution, from three separate experiments is presented in the tables.

One needs to be critically aware that, in the human airways, the porosome, in addition to appropriate secretion of mucus, is likely involved in the secretion of vesicular bodies that maintain normal lung physiology and function [54,55]. Our recent studies [56] on brain neurons and insulin-secreting Min6 cells suggest that in addition to the docking, fusion and content release, the porosome may also participates in the release of nanoscale vesicles. Electron micrographs (EM) show that in addition to neurotransmitter containing synaptic vesicles and insulin containing granules, multi-vesicular bodies and exosome-like vesicles are also released by the porosome complex in these cells. In addition to EM micrographs, this was further confirmed using porosome protein knockout studies on Min6 cells [56]. The porosome-associated calcium transporting ATPase 1 [SPCA1] encoded by the ATP2C1 gene, when knocked-out in Min6 cells using CRISPR, a loss of both glucose-stimulated insulin secretion and exosome release was observed in the ATP2C1 knockout Min6 cells [56]. These results further support that altered porosomes in CF need to be addressed via the reconstitution of normal functional mucin-secreting porosomes, since CFTR mutation may alter the secretion of such vesicular bodies and impact tissue pathophysiology. The WT-CFTR HBE cell line could be used as a source for harvesting normal functional mucin-secreting porosomes for porosome-reconstitution therapy in the lung, and could easily be scaled up as required. In addition to the WT-CFTR HBE cells grown in media, the differentiated 3D ALI cultures could be used. Similarly, ALI certified normal human bronchial epithelial cells using an established ***rotating bioreactor for scalable culture and differentiation of the human respiratory epithelium*** [57] could also be utilized.

Functional reconstitution of the porosome in live cells holds great promise for future therapeutic applications. Cellular secretory defects resulting from impaired porosome functions could be overcome by reconstituting isolated porosomes from healthy tissue. Since the porosome is a nanoscale cellular structure present in all secretory cells, its reconstitution is unlikely to elicit an immune response and therefore in various secretory diseases could be used for therapeutic applications. For example, porosome reconstitution could be advantageous in optimizing the secretory capability of various tissue transplants. A major issue in treating Type 1 Diabetes (T1D), using beta cells derived from induced pluripotent stem cells (iPSC), is the inability of these beta cells to optimally secrete insulin in response to a glucose challenge. Consequently, a large number of such iPSC-derived beta cells is required for a transplant in the treatment of T1D, which poses a significant clinical problem. This problem can now be overcome by reconstituting insulin-secreting porosomes [28] into the cell plasma membrane of iPSC-derived beta cells prior to their transplant in patients. Reconstitution will restore normal insulin secretion capability to these iPSC-derived beta cells.

## Ongoing Studies

Current ongoing studies involve **a)** porosome dose optimization to obtain a 100% normal mucin secretion in CF, **b)** optimization of porosome reconstitution efficiency, **c)** determine porosome reconstitution viability to determine treatment frequency, and **d)** toxicity and immune-compatibility studies to evaluate cellular responses such as cell surface marker expression and localization such as CD86 (pro-inflammatory) and CD206 (anti-inflammatory). CD86 is present in antigen presenting cells which provides signal for T cell activation. CD86 is a pro inflammation M1 lineage receptor. CD206 is a M2 subtype pattern recognition receptor which mediates phagocytosis [58].

It is important to recognize that the human airway epithelium, being the primary target for all inhaled air-borne substances, needs to have an appropriate model when evaluating the potency, efficacy and toxicity of new lung-targeted drugs and therapies. In porosome-reconstitution therapy, the nano-sized porosome complex is retained at the cell plasma membrane. Recent developments in advanced 3D ALI bronchial epithelia organoids, including human primary epithelial cells and induced human pluripotent stem cells for generation of lung epithelium, are among the valid methods that may now be suitable to replace animal models for inhaled toxicity studies and lung therapies. Toxicity and efficacy studies on lung-based drug delivery however, is a different matter.

## Contributions

The idea for the use of porosome-reconstitution therapy was developed at Porosome Therapeutics, Inc. The research design was developed by B.P.J. and conducted by W-J.C. MS/MS studies were performed as paid contract by the Wayne State University Proteomics Core. B.P.J. wrote the paper. Both authors participated in critical reading and discussion of the manuscript.

## Acknowledgements

Work presented in this article was supported by the Viron Molecular Medicine Institute and Porosome Therapeutics, Inc., Boston, MA. We thank Prof. Daniel A. Walz for his critical reading of the manuscript and his valuable suggestions.

